# Decision-making strategies of two related fish species diverge under increased perceptual load

**DOI:** 10.1101/2025.03.07.641999

**Authors:** Maëlan Tomasek, Boyd Dunster, Zoë Goverts, Dylan Naceur, Alex Jordan, Valérie Dufour

**Affiliations:** LAboratoire de Psychologie Sociale et COgnitive, UMR6024, CNRS, UCA, 63000 Clermont-Ferrand, France; Behavioural Evolution Research Group, Max Planck Institute of Animal Behaviour, 78467 Konstanz, Germany

**Keywords:** preferences, multi-attribute decision-making, perceptual load, comparative cognition, framing effects, attention

## Abstract

Three main mechanisms have been proposed to explain divergent decisions between species faced with the same options: differences in sensory abilities, attentional capacities, or cognitive evaluations. While these mechanisms have been well-established in controlled settings, there is limited empirical evidence regarding the ways in which different species make decisions under varying perceptual loads. Here we investigated the decision-making processes of two closely related cichlid species, *Aulonocranus dewindti* and *Cyathopharynx furcifer*, in Lake Tanganyika. Males of these species construct sand bowers from which they remove foreign objects, and we examined their preferences when presented with objects varying in three attributes: colour, size, and shape. We show that both species exhibited similar preferences under low perceptual load. Similarly, when the perceptual load was increased by increasing the number of options, both species’ choices were driven by attentional capture, and an oddity effect was observed. However, when the perceptual load was increased by adding more attributes (feature conjunctions), the species diverged in their decision-making. One species, *A. dewindti,* evaluated the options in an absolute manner, considering all available attributes, as evidenced by its insensitivity to decoy effects. The other species, *C. furcifer* either showed attentional capture or evaluated the options in a comparative manner. We discuss the potential reasons behind these differences and their implications for the evolution of decision-making mechanisms.

## 1. Introduction

Among the many decisions they make during their day, animals often express clear preferences in their behaviour. For example, flower-visiting insects display preferences in flower colours, sizes or patterns ^1^, and bowerbirds choose to integrate or remove objects from their bowers based on colours ^2,3^. When several options are available, deciding which option to select can be more or less complex, depending on the characteristics of these options. Imagine yourself at a restaurant. The waiter is coming and you have a few seconds to make a decision. Fortunately, the menu is composed of two dishes, both of the same price. In this case, you will easily make a decision based on your food preferences. Choices between few options differing in only one feature (here, the nature of the dishes) are of low perceptual load and can be easily processed ^4^. In some situations, however, the perceptual load can increase and require more attentional capacity ^4^. This is the case when the menu presents more than two dishes. This is also the case when the menu is composed of two dishes that are of different prices with options that are conflicting: your preferred dish is more expensive.

In each case, the chooser’s attentional capacity can be overloaded, and the information available may be entirely processed. When the number of options increases, decisions can be based on phenomena such as “attentional capture” also referred to as selective bottom-up attention, where the most salient option is chosen ^5,6^. For instance, predators hunting a group of animals are often attracted to prey which are most noticeable among the group: this is called the oddity effect ^7,8^. In the second case, the perceptual load is increased because individuals have to take into account several features in each option, i.e. feature conjunctions ^4,9–11^. Animals often face such multi-attribute decisions, for instance in mate choice in the context of sexual selection ^12^. If attentional capacity allows it, the options are cognitively evaluated. This can be done in an absolute manner: individuals assign an inherent value to each option independently of the others. According to some authors, such computations could be the most cognitively demanding ^13^. Animals can also evaluate the options in a comparative manner where the value of each option will be dependent of the other available options ^12^. One example of such strategy is to rank the attributes of the options and base one’s decision only on the important ones. For instance, *Macroglossum* hawkmoths give more importance to flower size than colour, and more importance to colour than pattern ^14^. *Vanessia* butterflies prioritize flower colour over scent ^15^. However, even when using simple strategies, multi-attribute decision-making is often complex and can depend on context or experience ^1,16^.

One way to assess whether an animal uses an absolute or comparative evaluation of options in multi-attribute decision-making is to investigate a particular case of framing effects (cognitive biases in which the presentation of the options influences the decisions ^17^), the decoy effect, where decisions can be influenced by irrelevant alternatives. Let’s suppose two options, A and B, that are equally preferred. The introduction of a third option C (the “decoy”), which is less preferred than B (the “competitor” option) on one attribute and less preferred than A (the “target” option) on two attributes, should not influence the preferences between A and B in an absolute evaluation of the options. The “constant ratio rule” states that the relative proportion of choices between A and B should remain equal when C is added ^18^. However, if the animal uses a comparative evaluation of the options, the addition of C can modify the preference relationship between A and B, leading to a decoy effect (see ^1^ for a review).

Interspecific differences in information processing during a choice can be explained by three mechanisms: divergence in sensory abilities, attentional capacities, and cognitive evaluation. In closely related species with similar sensory skills, providing choice situations with high perceptual load is a way to disentangle the relative role of attentional versus cognitive processes in choices. Indeed, different species could have evolved different attentional capacities, enabling some to process more information than others. Likewise, in identical situations, different species might cognitively evaluate the options in a different manner, using divergent absolute or comparative strategies. The adaptive radiation of Tanganyikan cichlids is a prime system to investigate cognitive divergences throughout evolutionary history, as the lake is home to more than 250 closely related species displaying a wide variety of life history traits ^19^. The present study investigates decision-making in the wild in two closely related species of Tanganyikan cichlid fish: *Aulonocranus dewindti* and *Cyathopharynx furcifer* (**Figure 1A**). These two species are phylogenetically close as their last common ancestor is estimated at 2 million years ago ^19^. Both species live in the same habitat and are organized in a lekking system in which males build sand bowers to attract females ^20,21^. They maintain their bowers clean and both species display the same decision rule when removing objects from their bower: if a snail shell and a stone are put inside the bower, they generally prefer to remove the snail shell first ^21,22^. Interestingly, when comparing species in a preference reversal test (by bringing the shell back in the bower if it was removed first), we found cognitive differences in their performance with *A. dewindti* showing more inhibitory control in selecting the stone first than in *C. furcifer* ^21^. Here, we investigate decisions in these species that could hint at differences in sensory, attentional, or cognitive mechanisms. First, we explore the species’ preferences for three attributes of objects (colour, size, and shape) in binary choice tasks which are of low perceptual load. We then increase the perceptual load of the choices by presenting two situations. First, we add more options to the choices to investigate their susceptibility to an oddity effect. Then, we investigate their ability to process conflicting attributes in multi-attribute choices and, if relevant, substantiate an absolute or comparative evaluation of the options by testing their susceptibility to a decoy effect. Given the relatively small amount of time between the divergence of the two species, we do not expect large differences in decision-making mechanisms. However, considering the remarkable evolutionary rapidity of the Tanganyikan cichlid radiation, it could still be possible to find subtle differences between both species.

**Figure 1:**
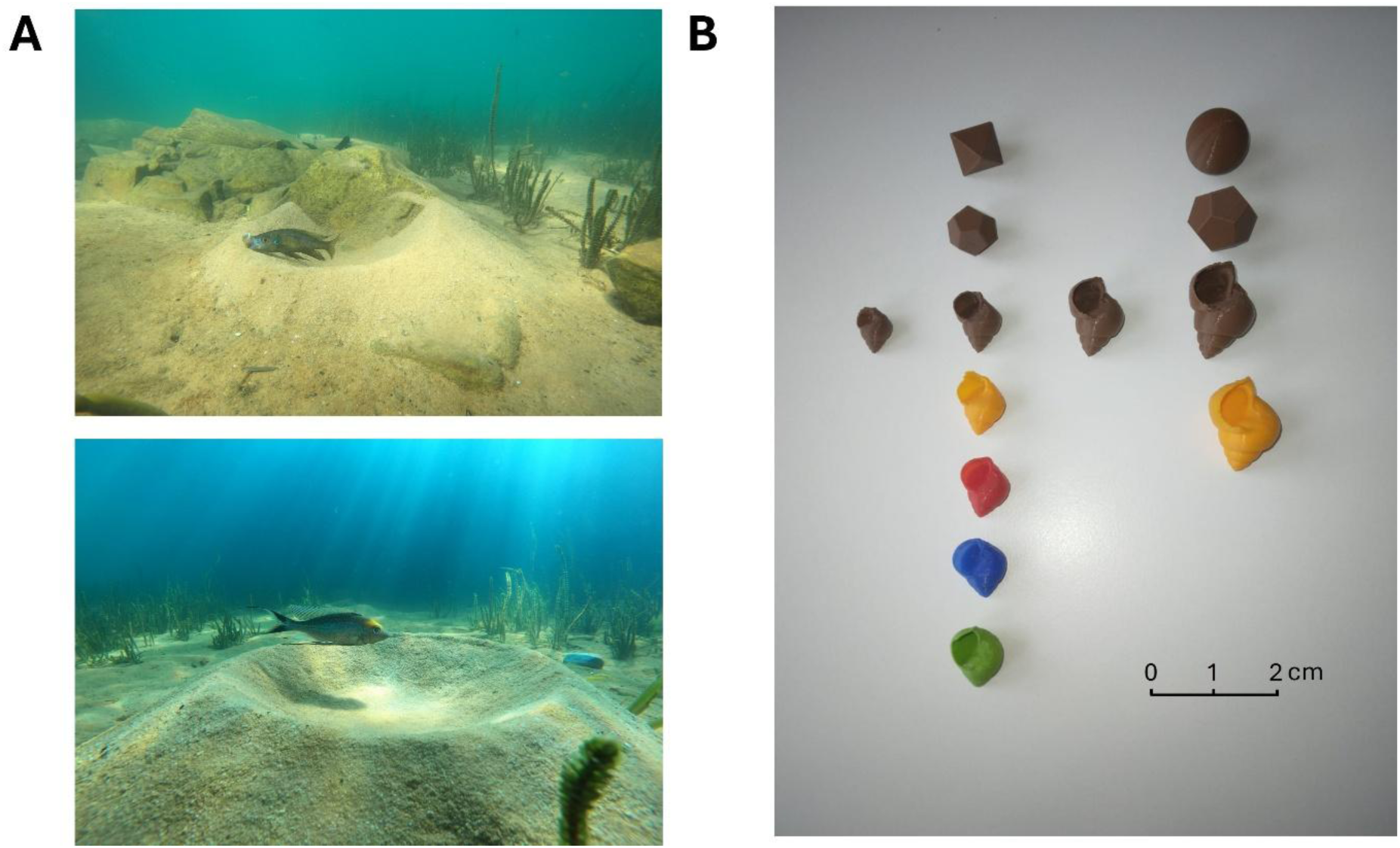
Species and materials. **A.** Study species: *Aulonocranus dewindti* (above) and *Cyathopharynx furcifer* (below). **B.** 3D-printed objects used in the experiments.

## 2. Results

Adult male *Aulonocranus dewindti* and *Cyathopharynx furcifer* were found along the south-eastern Zambian shore of Lake Tanganyika at depths between 3m and 6m. All experiments were conducted while SCUBA-diving. 3D-printed objects varying in colour (blue, orange, brown, red, and green), size (1.5cm, 2cm, 2.5cm, and 3cm), and shape (*Neothauma* shell, dodecahedron, octahedron, and sphere) were used for all experiments (**Figure 1B**).

### 2.1. Both species showed similar preferences for colour, size, and shape in situations of low perceptual load

We first assessed the baseline preferences in colour, size, and shape for both species in binary choice tasks of low perceptual load. We place two objects varying in only one attribute in the bower of the tested individual and we recorded which object was removed first. In total, we conducted 3322 trials in which we tested 15 binary combinations with a minimum of 6 individuals per species and 90 trials per combination. Due to time and practical constraints in the field, not all combinations could be tested.

Within each species, we looked at the number of individuals who had a significant preference to remove one object first over another. Both species presented preferences in size, shape, and colour (**Figure 2**, **Table I**). In both species, size affected choices similarly as individuals would always prefer to remove the bigger object first. They also presented preferences of shapes, preferring to remove shells over other objects, consistent with previous observations that both species preferred to remove snail shells over stones^21^. Finally, both species presented preferences in colour. The two species showed similar patterns regarding the direction and strength of these preferences. Only two combinations showed differences in strength: *C. furcifer*’s preferences to remove red over blue and green over blue were stronger than *A. dewindti*’s (generalised linear mixed-effect models, estimates = 0.87 ± 0.40 and 0.94 ± 0.44 respectively, p-values = 0.03 and 0.03 respectively, **Table I**).

**Figure 2:**
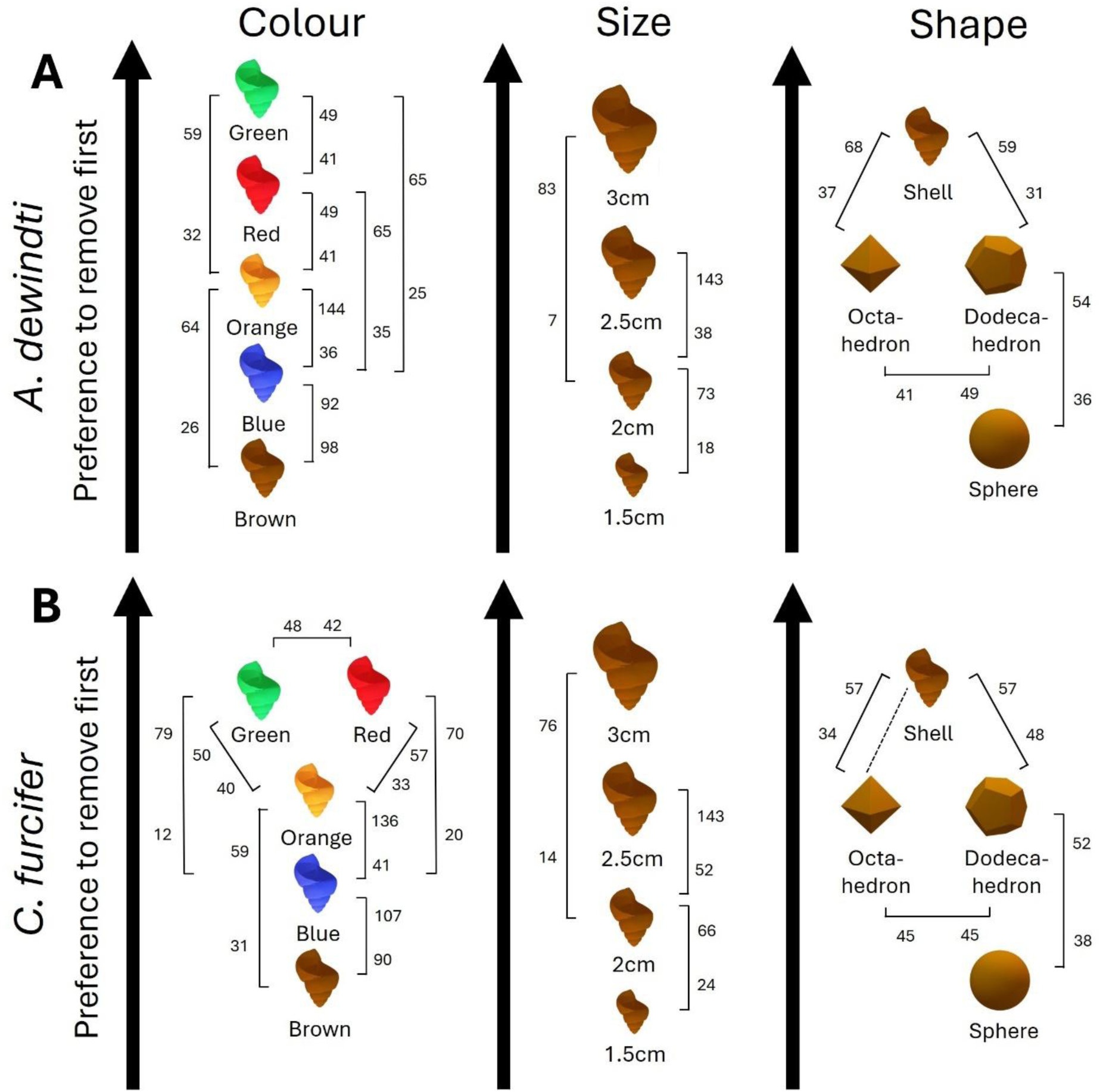
Baseline preference patterns inferred from binary choices in. **A.** *A. dewindti* and **B.** *C. furcifer*. The relative positions of the objects are based on the number of individuals presenting a significant preference (as soon as one individual tested significantly preferred removing one object first over the other, this object is considered as more preferred) and the numbers indicated represent the total choices of each option for every binary choice in the whole population. The dotted line between the octahedron and the shell for *C. furcifer* means that no individual showed any significant preference for one option and the options should be considered at the same level.

**Table I.** Results of the single-attribute binary choices. . Are indicated the number of total first removals in the whole population for each binary choice, the number of individuals presenting a significant preference for removing one option first or no preference (GLMM Object removed first ∼ Subject + (1|Session/Trial), binomial distribution), and the results of the interspecific differences in decision patterns (GLMM Object removed first ∼ Species + (1|Subject) + (1|Session/Trial), binomial distribution).

| SINGLE-ATTRIBUTE DECISION-MAKING |  |  |  |  |  |  |  |  |  |  |
| --- | --- | --- | --- | --- | --- | --- | --- | --- | --- | --- |
| Colour binary choice (Shells, 2cm) |  |  |  |  |  |  |  |  |  |  |
| Colour A | Colour B | Species | Number of choices |  | Number of individuals |  |  | Interspecific difference |  |  |
|  |  |  | A | B | Preference for A | No preference | Preference for B | Estimate | Standard error | p-value |
| Blue | Brown | Dewindti | 92 | 98 | 1 | 10 | 0 | -0.23 | 0.37 | 0,54 |
|  |  | Furcifer | 107 | 90 | 3 | 9 | 1 |  |  |  |
| Blue | Red | Dewindti | 35 | 65 | 0 | 6 | 1 | 0.87 | 0.40 | 0,03 * |
|  |  | Furcifer | 20 | 70 | 0 | 5 | 1 |  |  |  |
| Blue | Green | Dewindti | 25 | 65 | 0 | 4 | 2 | 0.94 | 0.44 | 0,03 * |
|  |  | Furcifer | 12 | 79 | 0 | 0 | 6 |  |  |  |
| Blue | Orange | Dewindti | 36 | 144 | 0 | 2 | 4 | -0.29 | 0.52 | 0,59 |
|  |  | Furcifer | 41 | 136 | 0 | 1 | 5 |  |  |  |
| Brown | Orange | Dewindti | 26 | 64 | 0 | 3 | 3 | -0.29 | 0.50 | 0,56 |
|  |  | Furcifer | 59 | 31 | 0 | 5 | 1 |  |  |  |
| Green | Orange | Dewindti | 59 | 32 | 1 | 5 | 0 | 0.38 | 0.39 | 0,34 |
|  |  | Furcifer | 50 | 40 | 1 | 5 | 0 |  |  |  |
| Orange | Red | Dewindti | 41 | 49 | 1 | 5 | 0 | 0.38 | 0.38 | 0,32 |
|  |  | Furcifer | 33 | 57 | 0 | 5 | 1 |  |  |  |
| Green | Red | Dewindti | 49 | 41 | 1 | 5 | 0 | 0.07 | 0.67 | 0,92 |
|  |  | Furcifer | 48 | 42 | 2 | 2 | 2 |  |  |  |
| Size binary choices (Shells, brown) |  |  |  |  |  |  |  |  |  |  |
| Size A | Size B | Species | Number of choices |  | Number of individuals |  |  | Interspecific difference |  |  |
|  |  |  | A | B | Preference for A | No preference | Preference for B | Estimate | Standard error | p-value |
| 1.5cm | 2cm | Dewindti | 18 | 73 | 0 | 3 | 3 | -0.39 | 0.42 | 0.35 |
|  |  | Furcifer | 24 | 66 | 0 | 3 | 3 |  |  |  |
|  |  | Furcifer | 52 | 143 | 0 | 1 | 6 |  |  |  |
| 2cm | 3cm | Dewindti | 7 | 83 | 0 | 0 | 6 | -0.81 | 0.50 | 0.11 |
|  |  | Furcifer | 14 | 76 | 0 | 1 | 5 |  |  |  |
| Shape binary choices (Brown object, 2cm or 3cm for Dodecahedron - Sphere) |  |  |  |  |  |  |  |  |  |  |
| Shape A | Shape B | Species | Number of choices |  | Number of individuals |  |  | Interspecific difference |  |  |
|  |  |  | A | B | Preference for A | No preference | Preference for B | Estimate | Standard error | p-value |
| Dodecahedron | Shell | Dewindti | 59 | 31 | 0 | 5 | 1 | -0.53 | 0.35 | 0.13 |
|  |  | Furcifer | 57 | 48 | 0 | 5 | 1 |  |  |  |
| Octahedron | Shell | Dewindti | 68 | 37 | 0 | 6 | 1 | -0.09 | 0.31 | 0.78 |
|  |  | Furcifer | 57 | 34 | 0 | 6 | 0 |  |  |  |
| Dodecahedron | Octahedron | Dewindti | 49 | 41 | 0 | 6 | 0 | 0.19 | 0.31 | 0.54 |
|  |  | Furcifer | 45 | 45 | 0 | 6 | 0 |  |  |  |
| Octahedron | Sphere | Dewindti | 54 | 36 | 1 | 5 | 0 | 0.04 | 0.43 | 0.93 |
|  |  | Furcifer | 52 | 38 | 1 | 6 | 0 |  |  |  |

### 2.2. Both species show attentional capture in a situation of high perceptual load with increased number of options

To investigate decision processes in situations of higher perceptual load, we increased the number of options to detect a possible oddity effect. We used two colours for which neither species had shown a preference: blue and brown. A trial consisted of the experimenter placing four 3D-printed 2cm *Neothauma* shells in the bower simultaneously: three shells were of the same colour and one shell was of the other colour (the odd shell). The trial ended once the fish had removed all shells. In both species, for both colours of odd shell (blue or brown), we conducted 15 trials per individual with 12 individuals, for a total of 180 trials for each colour of odd shell per species (720 trials in total). Almost all individuals (20 out of 24) had not shown a significant preference for one colour over the other in the binary choice task. If fish possess the attentional capacity to consider all options equally, then they should stick to their initial lack of preference and remove the odd shell equally often between all ranks of removal (25% of the trials removed first, 25% removed second etc.). If not, they can be subject to attentional capture, being attracted to the odd shell and showing an oddity effect where fish remove the odd shell first more often than 25% of the trials.

Both species showed such an oddity effect: fish preferred to remove the odd shell first more than other ranks of removal (**Figure 3**, ANOVA χ^2^ = 99.18, p<0.001). In this situation of high perceptual load, fish decisions were based on attentional capture.

**Figure 3:**
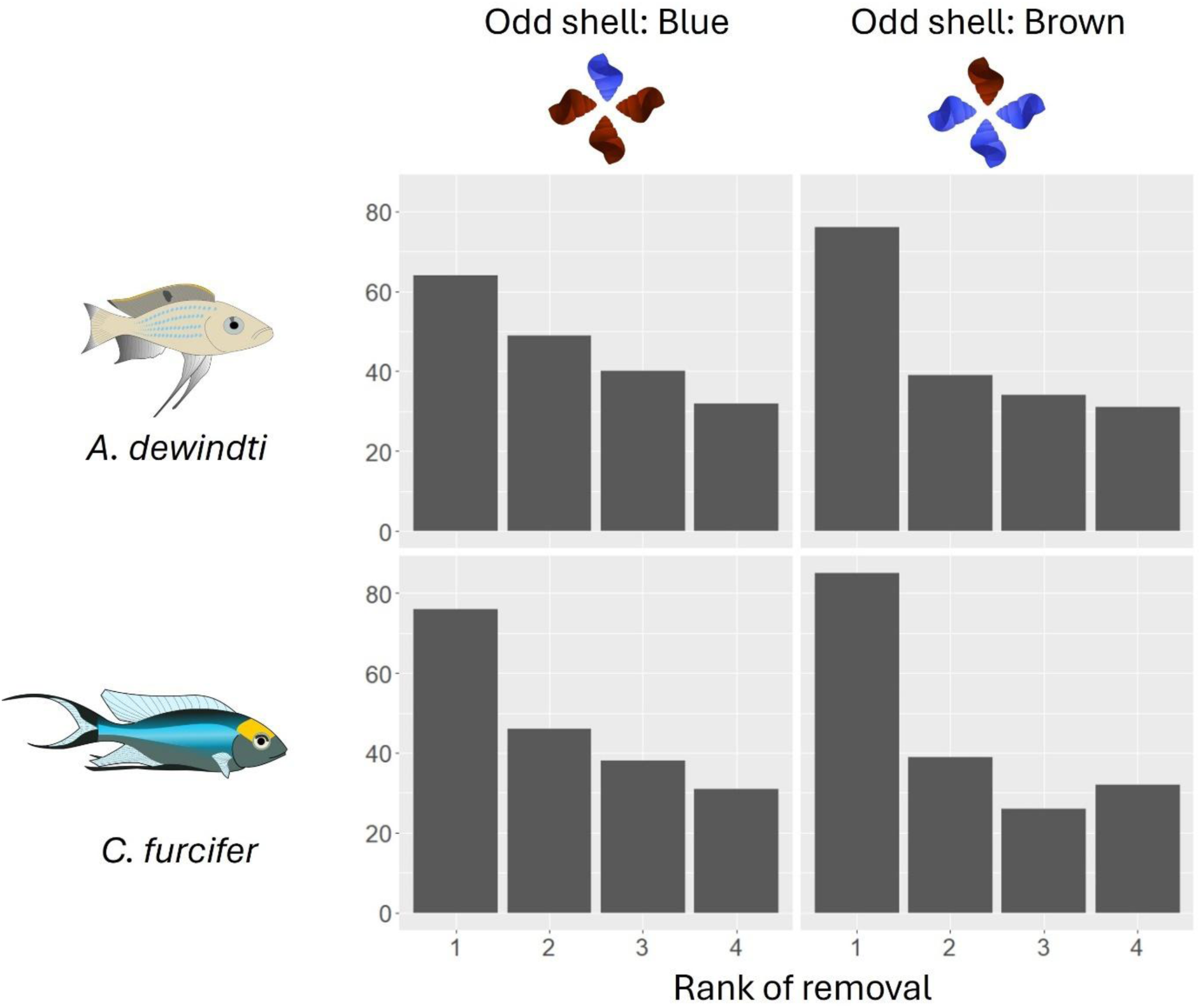
Decisions under high perceptual load (increased number of options, oddity effect). Counts of the different ranks of removal (odd shell removed first, second, third, or last) at the population-level. There is a significant effect of the rank in the whole population: both species significantly remove the odd shell first (GLMM Total of removals ∼ Rank * Species * Colour of the odd shell + (1|Subject) with a Poisson distribution, (ANOVA χ^2^ = 99.18, p<0.001, n = 180 trials per species per odd shell colour).

### 2.3. The two species showed different patterns of decision in situations of high perceptual load with increased number of attributes (feature conjunction)

Using the baseline preferences previously established, we then assessed decisions in binary choice tasks when the two objects presented two conflicting attributes (e.g., one object with a preferred size but unpreferred colour and the other object with unpreferred size but preferred colour). We investigated four combinations in *A. dewindti* and five in *C. furcifer*, for a total of 1470 trials in 17 *A. dewindti* and 12 *C. furcifer* (not all combinations were tested for all individuals).

When the options conflicted in size and shape, some individuals in both species showed a significant preference to remove a 2cm brown dodecahedron or a 2cm brown octahedron first (preferred size but unpreferred shapes) over a 1.5cm brown shell (preferred shape but unpreferred size). One individual out of six preferred to remove the dodecahedron or the octahedron first in all situations except for *C. furcifer* in which two individuals out of six significantly preferred to remove the dodecahedron first (**Figure** 4, Table **II**). Species did not differ significantly in their decisions patterns (generalised linear mixed-effect models, estimates = −0.30 ± 0.39 for dodecahedron and shell and 0.00 ± 0.31 for octahedron and shell, p-values = 0.44 and 1.0 respectively, **Table II**).

**Figure 4:**
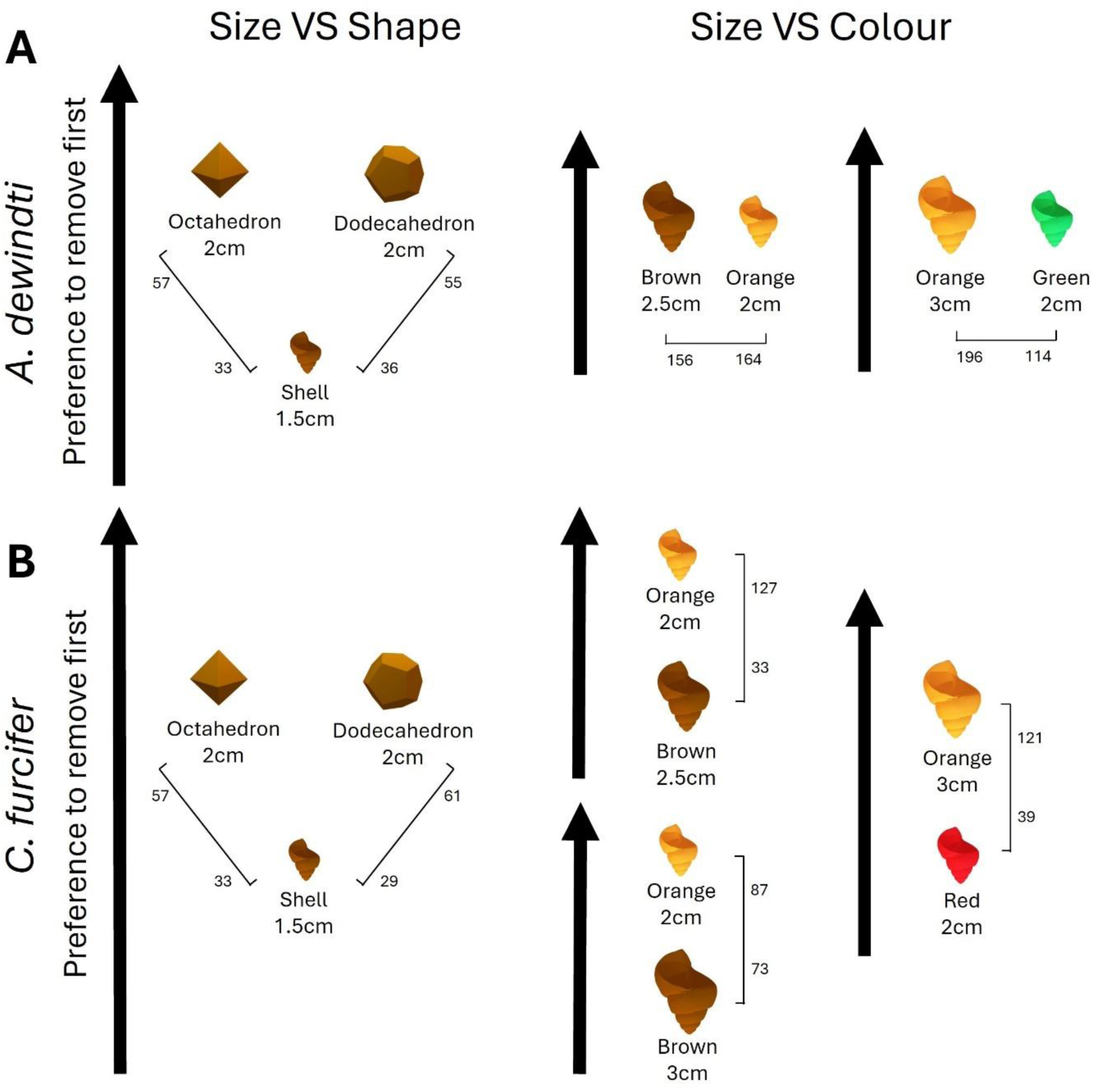
Decisions under binary choices of high perceptual load (feature conjunction) in. **A.** *A. dewindti* and **B.** *C. furcifer*. See Figure 2 for the meaning of the relative positions of the objects and the numbers indicated.

**Table II.** Results of the multi-attribute binary choices. See Table I for details. ^+^Model without the *C. furcifer* individual outlier presenting a significant preference for the red 2cm shell.

| MULTI-ATTRIBUTE DECISION-MAKING |  |  |  |  |  |  |  |  |  |  |
| --- | --- | --- | --- | --- | --- | --- | --- | --- | --- | --- |
| Shape VS Size (Brown objects) |  |  |  |  |  |  |  |  |  |  |
| Option A | Option B | Species | Number of choices |  | Number of individuals |  |  | Interspecific difference |  |  |
|  |  |  | A | B | Preference for A | No preference | Preference for B | Estimate | Standard error | p-value |
| Dodecahedron 2cm | Shell 1.5cm | Dewindti | 55 | 36 | 1 | 5 | 0 | -0.30 | 0.39 | 0.44 |
|  |  | Furcifer | 61 | 29 | 2 | 4 | 0 |  |  |  |
| Octahedron 2cm | Shell 1.5cm | Dewindti | 57 | 33 | 1 | 5 | 0 | 0.00 | 0.31 | 1.0 |
|  |  | Furcifer | 57 | 33 | 1 | 5 | 0 |  |  |  |
| Colour VS Size (Shells) |  |  |  |  |  |  |  |  |  |  |
| Option A | Option B | Species | Number of choices |  | Number of individuals |  |  | Interspecific difference |  |  |
|  |  |  | A | B | Preference for A | No preference | Preference for B | Estimate | Standard error | p-value |
| Brown 2.5cm | Orange 2cm | Dewindti | 156 | 164 | 1 | 13 | 2 | 1.54 | 0.46 | <0.001<br>*** |
|  |  | Furcifer | 33 | 127 | 0 | 2 | 6 |  |  |  |
| Brown 3cm | Orange 2cm | Furcifer | 73 | 87 | 0 | 5 | 3 | - | - | - |
| Green 2cm | Orange 3cm | Dewindti | 114 | 196 | 0 | 13 | 3 | -0.69 | 0.40 | 0.09 |
| Red 2cm | Orange 3cm | Furcifer | 39 | 121 | 1 | 1 | 6 | -1.06<sup>+</sup> | 0.34<sup>+</sup> | <0.01<br>** |

However, their decisions differed when the options conflicted in size and colour. For the first combination, *A. dewindti* did not show a preference between a 2.5cm brown shell (preferred size but unpreferred colour) and a 2cm orange shell (preferred colour but unpreferred size) (1 individual significantly preferred to remove the 2.5cm brown shell first and 2 the 2cm orange one, out of 16 individuals in total, **Table II**). On the contrary, *C. furcifer* showed a strong preference to remove the 2cm orange shell first over the 2.5cm brown shell (6 individuals out of 8 showed a significant preference, **Table II**). There was a significant difference in the decision patterns of the two species (generalised linear mixed-effect model, estimates = 1.54 ± 0.46, p < 0.001, **Table II**). Using the same individuals, we tested whether this preference was still present if the size of the shell was increased in *C. furcifer*. Although the preference vanished in some individuals, this species still showed a moderate preference to remove the 2cm orange shell first over the 3cm brown shell (3 individuals out of 8 showed a significant preference, **Table II**). In this context, *A. dewindti* therefore took both attributes into account, which led to no preference for any option, whereas *C. furcifer* focused on one attribute (colour).

For the second combination, we chose colours and sizes for which the two species had shown equivalent preference strengths in single-attribute decisions, and thus used a different colour for each species. *A. dewindti* showed a weak preference to remove a 3cm orange shell (preferred size but unpreferred colour) first over a 2cm green shell (preferred colour but unpreferred size) (4 individuals out of 16 showed a significant preference). *C. furcifer* showed a strong preference to remove a 3cm orange shell (preferred size but unpreferred colour) first over a 2cm red shell (preferred colour but unpreferred size) (6 individuals out of 8 showed a significant preference to remove the bigger orange shell first, but one individual showed a significant preference to remove the smaller red shell first, **Table II**). When this specific individual was removed from the analysis, there was a significant difference in the decision patterns of the two species (generalised linear mixed-effect model, estimates = −1.06 ± 0.34, p < 0.001, **Table II**). In this context, *A. dewindti* therefore took both attributes into account, which led to no preference for any option, whereas *C. furcifer* focused on one attribute (size).

Although both species presented the same preferences in single-attribute decisions, they acted differently in multi-attribute decisions when the colour and size of objects were in conflict: *C. furcifer* showed a preference for one of the two objects, while *A. dewindti* did not. This suggests that the latter species has the attentional capacity to take both attributes into account when making a decision and can evaluate the options in an absolute manner. *C. furcifer*, on the other hand, either does not have the attentional capacity to take both attributes into account, or if it does, evaluates the options in a comparative manner, prioritizing one attribute over the other.

### 2.4. *A. dewindti* uses absolute evaluation of the options in the feature-conjunction situation (no sensitivity to decoy effects)

To confirm whether *A. dewindti* uses absolute evaluation of the options in multi-attribute decision-making when size and colour are in conflict, we investigated whether the addition of a third irrelevant option (the “decoy” option) would modify the initial ratio of preferences between two options. We tested a decoy effect of size and of colour. In the size decoy experiment, we used a 2cm orange shell (“target” option) and a 2.5cm brown shell (“competitor” option) for which the fish showed no preference in experiment 2. We added a 1.5cm brown shell as a third option (“decoy” option) which, given the results of the baseline preferences, was less preferred than the competitor in one attribute (size) but less preferred than the target in both attributes (size and colour). In the colour decoy experiment, the “decoy” option was a 2cm blue shell which was added to a 3cm orange shell (“target” option) and a 2cm green shell (“competitor” option). This third option was less preferred than the competitor in one attribute (colour) but less preferred than the target option in both attributes (colour and size). If a decoy effect exists in this species, individuals should violate the constant ratio rule by modifying their initial relative preferences for the target and competitor options, which was 50/50 as they showed no preference for one option over the other in experiment 2. Note that we could not conduct these experiments with *C. furcifer* which already presented a strong preference for the target options in experiment 2. We only tested *A. dewindti* individuals which did not present an initial significant preference for the target options during the binary choice tasks. Thus, in total, we investigated a decoy effect of size in 14 individuals and of colour in 12 individuals, with a minimum of 20 trials per individual, for a total of 599 trials.

In the size decoy experiment, we found poor evidence for violation of the constant ratio rule in the population (**Figure 5A**). Only three individuals significantly changed their relative preferences proportions from experiment 2 to the decoy test (two for the target, one for the competitor). In the colour decoy experiment, we found no evidence for violation of the constant ratio rule in the population (**Figure 5B**). No individual switched its relative preferences from experiment 2 to the decoy test. Overall, *A. dewindti*’s decisions when size and colour were in conflict was not sensitive to a decoy effect which confirms that they evaluate these attributes in an absolute manner.

**Figure 5:**
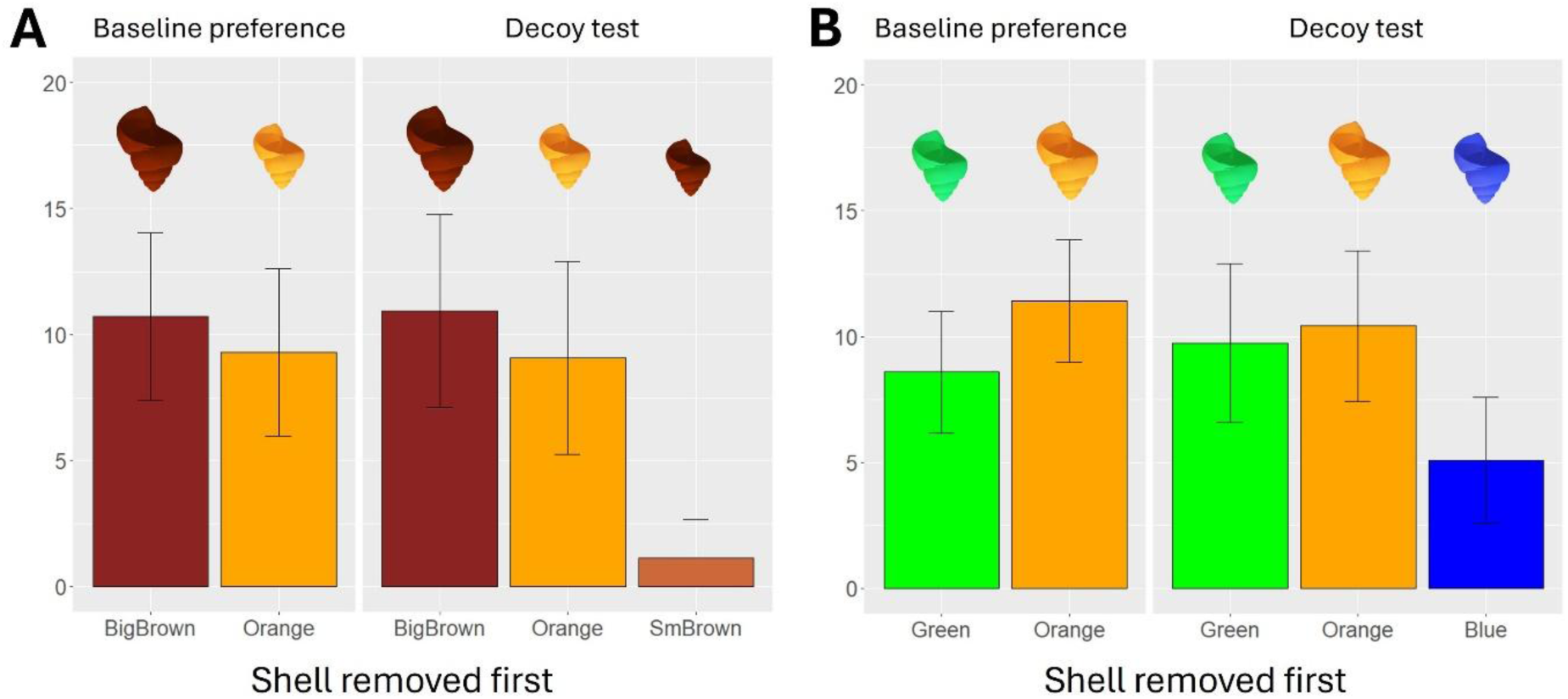
Decoy effect for size or colour in *A. dewindti*. **A.** Size decoy test (n = 14 individuals). **B.** Colour decoy test (n = 12 individuals). Left panels indicate the baseline preference between the target and competitor options in binary choice tasks and right panels show the results of the decoy tests. If *A. dewindti* was sensitive to a decoy effect, the ratio of preferences between target and competitor options should have been modified with the addition of the decoy option.

## 3. Discussion

In this study we found that two closely related cichlid species, *Aulonocranus dewindti* and *Cyathopharynx furcifer*, showed equal preference patterns for colour, size, and shape when removing objects from their bower in binary choices of low perceptual load. When the perceptual load was increased by presenting a set of options containing one odd coloured shell out of four, both species’ decisions were driven by attentional capture from the odd shell. When the perceptual load was increased by presenting multi-attribute choices, both species showed an equal preference to remove the bigger object first when the size and shape attributes of the options were in conflict. However, in combinations where colour and size were in conflict, decision patterns differed between both species. *C. furcifer* showed strong preferences to remove one of the two objects first, whereas *A. dewindti* individuals did not, suggesting that the latter species have the attentional capacity to consider all attributes in the decision and evaluate the options in an absolute manner. This was further substantiated by the lack of sensitivity to a decoy effect for size or colour in this species ^1,12^.

While decision patterns were similar between both species when the options differed in just one feature (colour, size, or shape), they showed different decisions in some contexts of multi-attribute choices. Several mechanisms could explain these differences. First, both species could differ in their sensory abilities, and decisions could for instance be underpinned by an impossibility to discriminate between objects instead of a real preference ^23^. However, both our species show similar decision patterns in single-attribute decisions, proving that they can discriminate equally well between all options, and there is no evidence for interspecific differences in sensory abilities. Another mechanism could be differences in attentional capacity, which can limit or expand access to information about the options in situations of high perceptual load ^24,25^. For instance, both species showed attentional capture in situations of high perceptual load due to an increase in the number of options. In situations of feature conjunction, *A. dewindti* take both attributes into account in their decisions but *C. furcifer* could have lower attentional capacity than *A. dewindti*, making them focus on the most salient feature of one of the options by attentional capture and impeding them from considering all attributes of both options ^5,6,26^. If, however, *C. furcifer* and *A. dewindti* have the same attentional capacity, then the interspecific differences can be explained by different cognitive evaluation processes. *A. dewindti* adopts absolute evaluation of the options, considering all attributes and evaluating the options independently. On the contrary, *C. furcifer* would adopt a comparative evaluation of the options, showing a strong preference for one option over the other. This species could thus use comparative mechanisms like a ranking of attributes ^25^. For instance, in both species, when size and shape were in conflict, size seemed to be more important than shape. In size/colour contexts however, there is no clear ranking of attributes in *C. furcifer*: In one context, colour would have been the most important attribute, while in the other it would have been size. Moreover, when the size of the bigger object was increased, some individuals cancelled their preference. If attribute ranking strategies are employed in this species, then they would be context dependent. Absolute evaluation, or “option-wise processing”, by considering all attributes is considered more cognitively demanding than “attribute-wise processing” ^1,13,27^. In a previous study, *A. dewindti* showed more inhibitory control abilities than *C. furcifer*, hinting for differences in degrees of cognitive flexibility between both species ^21^. This would be in line with greater cognitive capacities available in *A. dewindti* for decision-making.

Our results show that the evolution of decision-making mechanisms can be fast and subtle: in less than 2 million years, both species showed different decisions in some situations of high perceptual load. This difference can be explained either by difference in attentional capacities or in cognitive evaluation of the options. Further tests need to be conducted to specifically investigate differences in attentional capacities, testing for instance the perception of distractors in situations of varying perceptual loads ^4^. Adopting an adaptationist approach, we can hypothesize on the socio-ecological factors at the origin of this divergence ^28^. *A. dewindti* and *C. furcifer* do not differ in their social life history traits but mainly in two ecological traits: both are passive foragers but *C. furcifer* feeds on algae while *A. dewindti* feeds on plankton, and both build sand bowers but of different complexity (*C. furcifer* on flat sand or rock surfaces and *A. dewindti* among complex rock faces ^21,29^). It could be hypothesized that *A. dewindti* faces more complex challenges when building their bowers, having to take into account different features of the rocks. This could have selected for better attentional capacity and better cognitive processing of different attributes in high perceptual load situations. However, this could also just be a consequence of improved overall attentional or cognitive capacity selected in other non-decisional contexts. On the other hand, the slight differences in decision-making processes between our two species could also hint at non-adaptative mechanisms and be the consequence not of natural selection but of other non-directed evolutionary forces ^30^. Our findings encourage decision-making research to use similar paradigms in several species, even in relatively simple paradigms using differences in perceptual load. This could reveal differences in decision-making processes and hopefully shed light on their evolutionary mechanisms.

## 4. Methods

### 4.a. Experimental model and subject details

Adult male *Aulonocranus dewindti* and *Cyathopharynx furcifer* were found along the south-eastern Zambian shore of Lake Tanganyika (Isanga Bay, 8°37’24.7”S, 31°12’02.9”E) at depths between 3m and 6m. Of about 25 individuals of each species tested for candidacy, 18 *A. dewindti* and 16 *C. furcifer* were responsive and tested in at least one experiment. Responsive individuals were defined as such when they removed items from their bower within a maximum of three minutes. Note that they usually did so in a matter of seconds. Individuals could be recognized from day to day because bower-holding males hold the same bower for an extended period of time ^31^. They could also be further distinguished based on physical differences (size, body markings, and unvarying colour hues). Experiments were conducted while scuba diving and took place between 06:00 and 11:00 AM between April 6^h^ and May 13^th^, 2024. An external clinical examination of all potential candidates was conducted by a trained veterinarian (MT) and revealed no obvious health issue.

### 4.b. Binary choice tasks

Binary choice tasks were used to assess the preference of the fish to remove one object first when presented with two objects in their bower. A trial consisted of the experimenter placing two 3D-printed objects (polylactic acid) inside the bower of a fish and waiting until both were removed. Objects were placed in the center of the bower, a few centimeters apart, in a standardized position. To avoid location biases, their relative positions were alternated at each trial. A session consisted of five consecutive trials. If several sessions were conducted with the same individual within the same dive, the experimenter swam away for at least three minutes between sessions. For each binary combination, the minimal experimental structure consisted of three sessions of five trials in six individuals per species, for a total of 90 trials per species. Deviations from this basic structure occurred 14 times out of 39 experiments in total, either when more than six individuals were tested (to obtain baseline preferences for further experiments or because some individuals became unreactive during the experiment) or when more trials were conducted per individual.

In situations of low perceptual load, objects varied in only one attribute: colour, size, or shape. For colour, we used 3D-printed *Neothauma* shells of 2cm length varying in colour (blue, orange, brown, red, and green). For size, we used 3D-printed *Neothauma* brown shells varying in length (1.5cm, 2cm, 2.5cm, and 3cm). For shape, we used 3D-printed brown objects of either 2cm or 3cm length varying in shape (*Neothauma* shell, dodecahedron, octahedron, and sphere). Note that due to time and practical constraints in the field (not all objects could be fabricated and no 3D-printer was accessible), not all combinations could be tested. In situations of high perceptual load (feature conjunction), objects varied in two attributes that were in conflict (e.g. one object with a preferred size but unpreferred colour and the other object with unpreferred size but preferred colour).

Each session was videorecorded and we scored in each trial the first object removed from the bower (defined as the first object crossing the rim of the bower). If an individual clearly showed an intent of removing an object first (grabbing and displacing it) but let this one slip from its mouth before it crossed the rim of the bower (which happened in 2% of all trials, 84 times out of 4792 trials), we counted this attempt as the first object removed.

### 4.c. Oddity effect: high perceptual load with increased number of options

In both species, we investigated whether increasing the perceptual load by adding more options would influence fish decisions. More precisely, we presented a set of three equal options and one odd option. We used two colours for which most individuals in both species had not shown a preference: blue and brown (only 4 out of the 24 tested individuals had shown a significant preference for one of the two colours in baseline preference assessments). A trial consisted of the experimenter placing four 3D-printed 2cm *Neothauma* shells in the bower simultaneously: three shells were of the same colour, and one shell was of the other colour (the odd shell). The trial ended once the fish had removed all shells. The shells were placed simultaneously in the bower in a standardized way with the tips facing each other, and the position of the odd shell was alternated at each trial. In both species, for both colours of odd shell (blue or brown), we conducted three sessions of five trials per individual with 12 individuals, for a total of 180 trials for each colour of odd shell per species. Each session was video recorded, and we scored the rank of removal of the odd shell (whether removed first, second, third, or last) for each trial.

### 4.d. Decoy effect test

All initial ratios of preferences were previously assessed during the multi-attribute decision-making choices. We kept only individuals which did not present an initial significant preference for the target options. Thus, in total, we investigated the decoy effect of size in 14 individuals and of colour in 12 individuals. A trial consisted of the experimenter placing the three options in the bower and waiting for the fish to remove all shells. All shells were placed in a standardized way with the tips facing each other, and the positions of the shells were alternated at each trial. A session consisted of five trials. To enable a thorough comparison with the baseline preference which consisted of 20 trials per individual, we conducted as many sessions of decoy tests as necessary to obtain 20 trials with either the target or the competitor option removed first. In a session of five trials, if the fish removed the decoy option first less than twice, the session was prolongated to have a total of five trials in which the fish removed either the target or the competitor option first (the maximum number of trials attained within such a session was nine). In a session of five trials, if the fish removed the decoy option first more than twice, the session was ended and another session was conducted later on to complete the missed trials (the maximum number of sessions that an individual did was five). Each session was videorecorded and we scored the removals of each option.

### 4.e. Quantification and statistical analysis

All statistical analyses were done using R (version 4.2.2).

#### 4.e.i. Binary choice tasks

In each species and for each combination of objects, we ran a generalized linear mixed-effect model (Object removed first ∼ Subject + (1|Session/Trial)) with a binomial family distribution (package “glmmTMB” in R; Brooks et al., 2017). For each generalized linear model, we analysed the residuals to explore the fit of the models (“DHARMa” package in R; Hartig, 2017). We thus obtained preferences for removing one object first at the individual-level.

To investigate species differences in the decision patterns (direction and/or strength), for each combination, we ran generalised linear mixed effect models (Object removed first ∼ Species + (1|Subject) + (1|Session/Trial)) with a binomial family distribution. Residuals were analysed as described above.

#### 4.e.ii. Oddity effect test

To investigate an oddity effect at the population level, we ran a generalized linear mixed effect model (Total of removals ∼ Rank * Species * Colour of the odd shell + (1|Subject)) with a Poisson family distribution. The variable “Rank” is the position in which the odd shell was removed (whether first, second, third, or last shell removed). Residuals of the model were analysed as described above. We then ran a type II ANOVA to analyse the contribution of each factor and their interactions (package “car” in R; Fox & Weisberg, 2019)).

To investigate an oddity effect at the individual level, for each subject, we conducted Pearson’s chi-square tests (package “stats” in R, R Core Team, 2022) to analyse whether the distribution of the ranks of removal of the odd shell was significantly different than a uniform distribution.

#### 4.e.iii. Decoy effect test

To investigate whether the ratios of preferences changed between the baseline preference experimental condition or the decoy effect experimental condition, we ran a generalised linear mixed effect model (Object removed first ∼ Experimental condition * Subject + (1|Session/Trial)) with a binomial family distribution. An analysis of the residuals and a type II ANOVA were conducted as described above. We then ran least-square means post-hoc tests to analyse whether specific individuals had modified their initial relative preference in the presence of the decoy (function “lsmeans”, “lsmeans” package in R; Lenth, 2016).

## 5. Acknowledgements

The authors gratefully acknowledge their collaborators at the Department of Fisheries in Mpulungu, Zambia, and at the University of Zambia. We gratefully thank Myriam Knöpfle and Paul Nührenberg, as well as all members of the Jordan Lab and LAPSCO for fruitful discussions.

## 6. Lead Contact

Further information and request for resources, including data and code, should be directed to and will be fulfilled by the lead contact, Maëlan Tomasek.

## 7. Declarations

### Ethical approval

We declare that our study is in full accordance with the ethical guidelines of our institutions and comply with the Zambian and European legislation on animal welfare. The experiments were conducted according to the Guidelines for the treatment of animals in behavioural research and teaching (https://doi.org/10.1016/S0003-3472(21)00389-4) and were conducted under Zambian Study Permit numbers C-24173000/4-24 and SP297185/4-22.

### Author’s contributions

M.T., V.D., and A.J. conceptualised the experiments. M.T., B.D., and Z.G. performed the experiments. M.T. analysed the data, performed the analyses, prepared the figures, and wrote the main manuscript text. D.N., V.D. and A.J. reviewed the manuscript. V.D. and A.J. contributed equally in supervising this work.

